# Loss of TIM4-Dependent Efferocytosis in Kupffer Cells Promotes Liver Fibrosis in Nonalcoholic Steatohepatitis

**DOI:** 10.1101/2024.01.30.578023

**Authors:** Hongxue Shi, Xiaobo Wang, Brennan Gerlach, Arif Yurdagul, Mary P. Moore, Faridoddin Mirshahi, Luisa Ronzoni, Arun J. Sanyal, Luca Valenti, Chyuan-Sheng Lin, Ira Tabas

**Affiliations:** Department of Medicine, Columbia University Irving Medical Center, New York, NY 10032, USA; Divsion of Gastroenterology, Hepatology and Nutrition, Department of Internal Medicine, Virginia Commonwealth University, Richmond, Virginia, USA; Precision Medicine Lab, Biological Resource Center, Department of Transfusion Medicine, Fondazione Ca’ Granda Ospedale Maggiore Policlinico Milano, Milano 20122, Italy; Department of Pathophysiology and Transplantation, Universitàdegli Studi di Milano, Milan 20122, Italy; Department of Pathology and Cell Biology, Columbia University Irving Medical Center, New York, NY 10032, USA; Department of Physiology and Cellular Biophysics, Columbia University Irving Medical Center, New York, NY 10032, USA

**Keywords:** nonalcoholic steatohepatitis (NASH), efferocytosis, TIM4, Kupffer cell, liver fibrosis

## Abstract

**Background and aims:** Hepatocyte apoptosis is a key feature of non-alcoholic steatohepatitis (NASH), but the fate of apoptotic hepatocytes in NASH is poorly understood. Herein we explore the hypothesis that impaired TIM4-mediated clearance of dead hepatocytes by liver macrophages (efferocytosis) is impaired in NASH and drives the progression to liver fibrosis.

**Methods:** Kupffer cell (KC)-TIM4 expression and efferocytosis were assayed in normal and NASH liver from humans and diet-induced NASH mice. The engulfment of human and mouse apoptotic hepatocytes by primary human and mouse liver KCs was assayed *ex vivo*. Causation was assessed in NASH mice using anti-TIM4 antibodies, KC-TIM4-knockout, or inducible KC-TIM4 expression, with analyses focused on efferocytosis of apoptotic hepatocytes by liver macrophages and liver fibrosis.

**Results:** In human and mouse NASH liver, apoptotic hepatocytes accumulated and was associated with the loss of the KC efferocytosis receptor TIM4. Anti-TIM4 inhibited the engulfment of apoptotic hepatocytes by primary human and mouse liver KCs *ex vivo*, and anti-TIM4 administration to early NASH mice worsened liver macrophage efferocytosis and accelerated the progression to fibrotic NASH. A similar result was obtained by genetically deleting TIM4 in KCs in NASH mice. Most importantly, genetic restoration of macrophage TIM4 in NASH mice enhanced the clearance of apoptotic hepatocytes by liver macrophages and decreased liver fibrosis.

**Conclusions:** The loss of macrophage TIM4 that occurs during NASH progression impairs the clearance of apoptotic hepatocytes by liver macrophages, which subsequently promotes the progression to fibrotic NASH. This pathogenic sequence of events can be prevented by restoring macrophage TIM4, suggesting that future therapeutic approaches designed to boost TIM4 expression in liver macrophages could represent a novel strategy to prevent fibrotic NASH progression.

**Lay summary:** Nonalcoholic steatohepatitis (NASH) is emerging as the leading cause of liver disease, but the processes leading to liver fibrosis in NASH, which determines clinical outcome, are incompletely understood. Our study provides evidence impaired clearance of dead liver cells by liver macrophages in NASH, which is due to loss of a macrophage receptor called TIM4, contributes to liver fibrosis. Knowledge of this process may suggest new ways to bolster the clearance of dead liver cells in NASH and thereby prevent the progression to liver fibrosis and subsequent liver disease.

## Introduction

The incidence of nonalcoholic fatty liver disease (NAFLD) is increasing to epidemic levels^1^. Among those with NAFLD, 10% to 30% will progress to nonalcoholic steatohepatitis (NASH), which is characterized by liver injury, hepatic steatosis, necroinflammation, and, most importantly, liver fibrosis due to activation of hepatic stellate cells (HSCs)^2,3^. NASH can lead to advanced liver cirrhosis, hepatocellular carcinoma, and liver failure, which significantly contributes to mortality in end-stage liver disease and a higher demand for liver transplantation. However, there are currently no FDA/EMA-approved drugs for NASH treatment^4^.

Previous studies have shown that multiple factors contribute to NASH pathogenesis, including fat accumulation in hepatocytes, oxidative stress, inflammasome activation, lipotoxicity, endoplasmic reticulum stress, and others^2,3^. These factors can cause hepatocyte death, notably apoptosis and necroptosis, which contributes to NASH progression, but the mechanisms linking hepatocyte death to HSC activation and NASH fibrosis remain poorly understood^5,6^. While there is evidence that apoptotic hepatocytes can release HSC activators^7,8^, clinical evidence that blocking apoptosis mitigates NASH is lacking. For example, the pan-caspase inhibitor emricasan failed to improve liver histology and paradoxically increased liver fibrosis in NASH patients^9^.

The accumulation of dead cells in disease settings implies that the process of dead cell clearance by phagocytic cells, or efferocytosis, is impaired^10,11^. Efferocytosis comprises a multi-step process in which "professional" phagocytes, notably macrophages, and other types of cells bind, internalize, and digest dead cells^10,11^. Macrophages bind apoptotic cells through efferocytosis receptors, such as TAM and TIM family members, leading to dead cell engulfment and degradation. The primary function of efferocytosis is to prevent dead cell accumulation and promote tissue resolution, and impaired efferocytosis contributes to many human chronic inflammatory diseases^10,11^. Little is known about dead cell clearance in NASH. Examples of the few reports in this area include reports that genetically induced knockout of macrophage TREM2, which can mediated apoptotic cell binding, impairs efferocytosis *ex vivo* and exacerbates experimental NASH in some studies but not others^12-15^ and that necroptotic hepatocyte clearance is impaired in NASH owing to upregulation of CD47 on necroptotic hepatocytes and its receptor SIRPα on NASH macrophages, leading to NASH progression^16^.

T cell immunoglobulin and mucin domain containing 4 (TIM4, encoded by *Timd4* gene), is a protein belonging to the TIM family and is primarily expressed in macrophages and dendritic cells^17^. TIM4 can function as an efferocytosis receptor and is expressed by liver resident Kupffer cells (KCs), and previous studies have shown that NASH liver macrophages have a marked decrease in TIM4 expression^18-21^. Although *in vitro* studies have shown that TIM4 functions in efferocytosis in isolated KCs^22-24^, its role in efferocytosis and liver fibrosis in NASH remain unknown.

Using analyses of human and mouse NASH liver, *ex vivo* and NASH liver efferocytosis assays, and causation studies in mouse NASH models, we provide evidence that the loss of macrophage TIM4 expression contributes to defective efferocytosis and apoptotic hepatocyte accumulation in NASH liver and fibrotic NASH progression. Most importantly, genetic restoration of TIM4 in macrophages in NASH improves both the clearance of apoptotic hepatocytes and dampens fibrotic NASH progression. These findings provide new mechanistic insight into NASH progression, which can inform future studies investigating new therapeutic strategies to block NASH fibrosis.

## Materials and Methods

### Animal studies

Male wildtype C57/BL6J mice (10-12 weeks old, #000664) were obtained from Jackson Laboratory (Bar Harbor, ME) and were allowed to adjust to the housing environment in the Columbia University Irving Medical Center for 1 week before starting experiments. The mice were fed a fructose-palmitate-cholesterol diet (FPC, Envigo, #TD. 160785 PWD) for 8 or 16 weeks to develop simple steatosis and fibrotic NASH, respectively^25^, or a high-fat choline-deficient L-amino acid-defined diet (HF-CDAA, Research Diets, #A06071302) for 4 or 10 weeks to induce early or advanced NASH, respectively^26^. Age-matched male mice were fed a control diet (PicoLab Rodent Diet 20, #5053). All the mice were randomly assigned to experimental groups by investigators, who were blinded in terms of group assignment. For the anti-TIM4 experiments, the mice were fed the HF-CDAA diet for 4 weeks, with anti-TIM4 antibody treatment (Bio-X-Cell, #BE0225, RRID:AB_2687708, 200 μg/mouse) or isotype control IgG treatment (Bio-X-Cell, #BE0225, #BE0089, RRID:AB 1107769, 200 μg/mouse) during weeks 3-4.

To generate *Timd4^fl/fl^* mice, a loxP-Neo-loxP (LNL) cassette was inserted in the intron upstream of the exon 2 of *Timd4* on a Bacterial Artificial Chromosome (BAC clone ID: RP23-646F6). The Neo cassette was removed by Cre recombinase to leave behind the first loxP site L83, and a Frt-Neo-Frt-loxP (FNFL) cassette was inserted in the intron downstream of the exon 2 of Timd4 gene. The Neo cassette was removed by Cre recombinase to leave behind the loxP site at L83. A gene targeting vector was constructed by retrieving the 2kb left homology arm (5’ to L83), the L83-exon2-FNFL cassette, and the 4kb right homology arm (end of FNFL cassette to 3’) into pMCS-DTA vector carrying a DTA (Diphtheria Toxin Alpha chain) negative selection marker. The FNFL cassette confers G418 resistance during gene targeting in FL19 (C57BL/6N) ES cells, and the DTA cassette provides an autonomous negative selection to reduce random integration during gene targeting. Several targeted ES cell clones were identified using PCR (primer sequences: F: ggaagcaattcccacctctatt; R: ctcccaaatcaagagaccagac) and injected into C57BL/6J blastocysts to generate chimeric mice. Male chimeras were bred to homozygous ACTB:FLPe female mice^27^ (Jackson Laboratory, #005703, C57BL/6J background) to transmit the floxed Timd4 allele. *Timd4^fl/fl^* mice were crossed with Clec4f-Cre mice^28^ (Jackson Laboratory, #033296) to generate KC-*Timd4* knockdown mice (KD); *Timd4^fl/fl^*mice were used as the wild-type control (WT).

To restore TIM4 expression in macrophages, we adopted a Tet-On system for doxycycline-inducible gene expression. A TRE (7 tetO+ minimal CMV promoter)—*Timd4*—bGHpA cassette was synthesized by GenScrip and injected into the pronuclei of fertilized C57BL/6J eggs to generate TRE-*Timd4* transgenic mice; PCR identified the transgenic founders using the TRE-*Timd4* primer sequences: F: ctcgtttagtgaaccgtcagat; R: CTGTCACCTCGATTGGTGTT. We crossed TRE-*Timd4* mice with *Cd68rtTA* mice (Jackson Laboratory, #032044, C57BL/6J background) to generate inducible macrophage TIM4 mice; TRE-*Timd4* mice or *Cd68rtTA* mice were used as controls. These mice, when they reached 10-12 weeks of age, were fed a HF-CDAA diet for 10 weeks, with doxycycline added to the drinking water (75 μg/ml) during weeks 6-10. Another group of these mice were fed the FPC diet for 16 weeks, with doxycycline (50 μg/ml) added to the drinking water (50 μg/ml) during weeks 9-16.

All mice were housed in standard cages at 22 °C in a 12-12-h light-dark cycle in a barrier facility, and experiments were conducted by following the Guidelines for the Care and Use of Laboratory Animals at Columbia University. Animal protocol AABL5573 for these studies was approved by the Institutional Animal Care and Use Committee at Columbia University.

### Human liver specimens (Table S1)

The de-identified FFPE human liver sections used for the in-situ efferocytosis assay in Fig. 1A were acquired from Virginia Commonwealth University and the Fondazione IRCCS Ca’ Granda Ospedale Policlinico, Universitàdegli Studi di Milano. Phenotypic and pathological characterizations were conducted by experienced medical physicians and pathologists associated with Virginia Commonwealth University and the Fondazione. The Fondazione patients underwent liver biopsy for suspected fibrotic NASH within a spontaneous observational study to investigate the role of genetic predisposition on the natural history of NAFLD. Patients gave informed consent at the time of recruitment, and their records were anonymized and de-identified. The de-identified human liver specimens used for Fig. 1C,D were acquired from the Liver Tissue Cell Distribution System at the University of Minnesota. The specimens were collected on the date of liver transplantation and preserved as frozen samples. Phenotypic and pathological characterizations were conducted by medical physicians and pathologists associated with the Liver Tissue Cell Distribution System. The diagnostic information for all samples is included in Table S1. All protocols were approved by the Institutional Review Board (IRB) at the Columbia University Irving Medical Center.

**Fig. 1.**
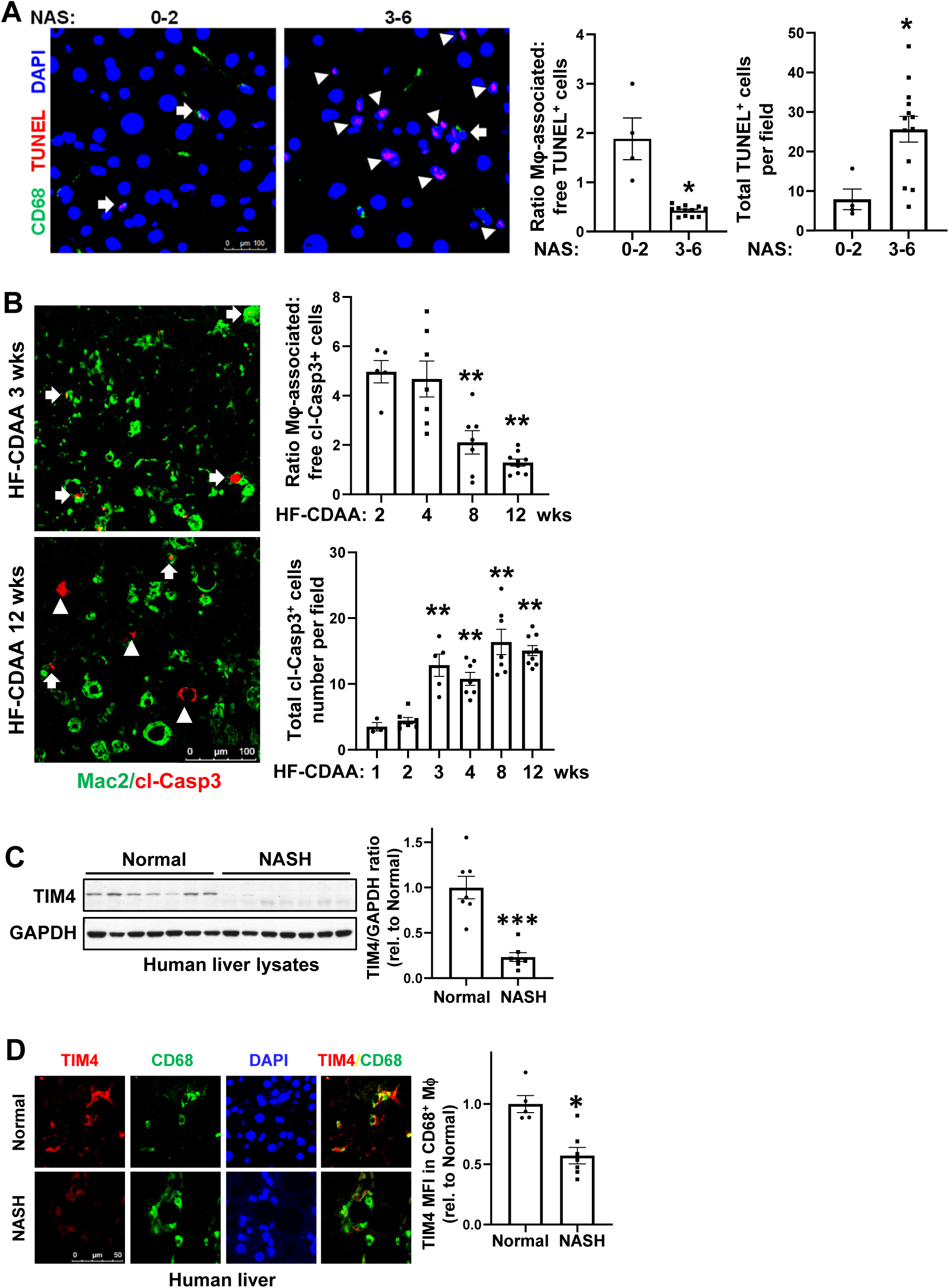
Efferocytosis by liver macrophages and the efferocytosis receptor TIM4 are decreased in human and experimental NASH. (**A**) Immunostaining of human liver sections from subjects with a NAS score of 0-2 or 3-6 using TUNEL (cell death) and anti-CD68 (macrophage, Mϕ). Arrows indicate macrophage-associated TUNEL^+^ cells, while arrowheads indicate macrophage-free TUNEL+ cells. Efferocytosis was determined by calculating the ratio of macrophage-associated: free TUNEL^+^ cells and total TUNEL^+^ cells per field (n = 4 in the NAS 0-2 group and 13 in the NAS 3-6 group). (**B**) Livers from mice fed the HF-CDAA diet for the indicated weeks were assayed for efferocytosis, using anti-cleaved caspase-3 (cl-Casp3) to mark apoptotic cells and anti-Mac2 to stain macrophages (n=3-9 mice per group. Examples of images at 3 and 12 weeks of diet feeding are shown. Bar, 100 μm. Arrows indicate macrophage-associated apoptotic cells, while arrowheads indicate macrophage free apoptotic cells. (**C**) Immunoblots of TIM4 in another cohort of human normal and NASH liver, with data quantification (n = 7/group). (**D**) Immunostaining of human normal and NASH liver sections from panel A using anti-TIM4 (red) and anti-CD68 (green). Bar, 50 μm. Data were quantified as TIM4 MFI in CD68^+^ macrophages relative to normal liver (n = 5 in normal group and n = 7 in NASH group). For immunofluorescence images, nuclei are stained with DAPI (blue). All data are means ± SEM. *p <0.05, **p <0.01, ***p <0.001 by Student’s t-test (panels A, C, and D) or one-way ANOVA (panel B).

### Liver primary cell isolation

Livers of 8-10 weeks old of male C57BL/6J mice were perfused with HBSS perfusion buffer (Corning, #21-022-CV) containing HEPES (Sigma, #H0887, 25 mM) and EDTA (Invitrogen, #15575-038, 0.5 mM), followed by HBSS digestion buffer (Gibco, #14025-092) containing collagenase D (4.4 U/mouse, Roche, #11088882001). The digested livers were gently ruptured to release cells, which were then centrifuged at 50 g for 1 min at 4 °C to pellet hepatocytes. Next, the hepatocytes were plated on collagen-coated cell culture dishes in DMEM/F12 medium with 10% FBS and 1% penicillin-streptomycin. The supernate fractions from the 50-g spin were used for KC isolation by centrifuging at 50 g for 1 min to remove residual hepatocytes. Then non-parenchymal cells were pelleted at 800 g for 7 min. KCs were isolated with anti-F4/80 beads (Miltenyi Biotec, #130-110-443) and cultured in DMEM medium containing 10% FBS and 1% penicillin-streptomycin. Primary human Kupffer cells (#HK1000.H15) and hepatocytes (#H1000.H15-3) were obtained from Sekisui XenoTech Company.

### Liver histology and plasma ALT activity measurement

Livers were fixed in 10% formalin for at least 24 hours at room temperature and then paraffin embedded and sectioned (8 μm thick) for histological analysis, including hematoxylin and eosin (HE) and Picrosirius Red staining (Polysciences, #24901) according to the manufacturer’s protocol. Plasma alanine transaminase (ALT) activity was measured using a commercial kit (Teco Diagnostics, #A526). Liver steatosis was calculated in HE-stained sections by measuring lipid droplet area using Image J.

### Liver immunohistochemistry

Liver paraffin sections were deparaffinized with xylene, hydrated, and then subjected to antigen retrieval using citrate sodium (Vector laboratory, #H-3300, 1:100 dilution) in a high-pressure cooker for 10 min. Next, the liver sections were incubated with 3% hydrogen peroxide (Sigma, #H1009) for 10 min at room temperature to block endogenous peroxidase activity, blocked with 5% donkey serum in PBS with 0.1% Triton X-100 (Sigma, #X100) for 1 hour at room temperature, and incubated at 4°C overnight with anti-mouse collagen 1a1 (Col1a1, Cell signaling technology, #72026S, RRID:AB_2904565, 1: 200 dilution), osteopontin (Opn, RD, #AF808, RRID:AB_2194992, 1:1000 dilution), and cytokeratin 19 (CK19, DSHB, TROMA-III, RRID:AB_2133570, 1:500 dilution) in PBS containing 1% donkey serum. The sections were then incubated with SignalStain^®^ Boost IHC detection reagent from Cell Signaling Technology (HRP, Rabbit, #8114; HRP, Rat, #72838; HRP, Goat, #63707), followed by color development using a DAB substrate kit (Cell signaling technology, #8059). Finally, the sections were counterstained with hematoxylin, dehydrated, and mounted. The images were captured using a Nikon microscope, and the data were analyzed using Image J software.

### Liver immunofluorescence staining and in-situ efferocytosis assay

Liver paraffin sections were deparaffinized, hydrated, and blocked with 5% donkey serum in PBS with 0.1% Triton X-100 for 1 hour at room temperature. For immunofluorescence experiments, human liver sections were incubated at 4 °C overnight with anti-human TIM4 (Cell signaling technology, #75484T, RRID:AB_2799871, 1:200 dilution), anti-CD68 (Agilent, #GA60961-2, RRID:AB_2661840, 1:500 dilution), and anti-cleaved caspase3 (Cell signaling technology, # 9661, RRID:AB_2341188, 1:100 dilution) in PBS containing 1% donkey serum. Mouse liver sections were incubated at 4 °C overnight with anti-mouse TIM4 (RD, #AF2826, RRID:AB_2201844, 1:200 dilution), anti-Clec4f (RD, #MAB2784, RRID:AB_2081338, 1:250 dilution), anti-Mac2 (Cedarlane, #CL8942AP, RRID:AB_10060357, 1:500 dilution), anti-F4/80 (Cell signaling technology, #70076s, RRID:AB_2799771, 1:200), anti-osteopontin (Opn, RD, #AF808, RRID:AB_2194992, 1:100 dilution), anti-α-smooth muscle actin (Sigma, #C6198, RRID:AB_476856, 1:100 dilution) and anti-cleaved caspase3 (Cell signaling technology, # 9661, RRID:AB_2341188, 1:100 dilution) in PBS containing 1% donkey serum. The sections were incubated with fluorescent dye-conjugated secondary antibodies (1:250 dilution) for 1 hour at room temperature, followed by nuclei staining with DAPI. For the in-situ efferocytosis assessment, human and mouse liver sections were incubated at 4 °C overnight with the pan-macrophage marker anti-Mac2 and either anti-cleaved caspase3 or In Situ Cell Death Detection Kit, TMR red kit (Sigma, # 12156792910) in PBS containing 1% donkey serum, followed by fluorescent dye-conjugated secondary antibodies administration. The ratio of macrophage-associated apoptotic cells to free apoptotic cells was quantified as a measure of efferocytosis, as described previously^29^. The images were captured using a Leica DMI 6000B fluorescence microscope, and the data were analyzed using Image J software.

### Immunoblotting

Liver protein was extracted using RIPA lysis buffer (Thermo, #89901) with a proteinase and phosphatase inhibitor cocktail (Thermo, #78445), followed by measurement of protein concentration using a BCA kit (Thermo, #23227). Each lane of 4-20% Tris-gels (Life technologies, EC60285) was loaded with 20 µg protein. After electrophoresis, the gels were transferred to nitrocellulose membranes (Bio-Rad, #1620115). The membranes were blocked with 5% non-fat milk in Tris-buffered saline with 0.1% Tween 20 (TBST) for 1 hour at room temperature and then incubated overnight at 4°C with antibodies recognizing human TIM4 (Cell signaling technology, #75484T, RRID:AB_2799871, 1:1000 dilution), mouse TIM4 (RD, #AF2826, RRID:AB_2201844, 1:1000 dilution), and GAPDH (Cell signaling technology, #3683, RRID:AB_1642205, 1:5000 dilution). The membranes were then incubated with HRP-conjugated second antibodies (Jackson ImmunoResearch, #711-035-152, RRID:AB_10015282, 1:5000 dilution, or #705-035-003, RRID:AB_2340390, 1:5000 dilution) for 1 hour at room temperature, and bands were detected with SuperSignal West Pico PLUS Chemiluminescent Substrate (Thermo, #34580).

### Quantitative RT-qPCR

Total liver mRNA was extracted using TRIzol Reagent (Thermo, #15596026) and purified using PureLink™ RNA mini kit (Thermo, #12183025) according to the manufacture’s instruction. The mRNA concentration was measured using a nano-drop spectrophotometer, and cDNA was synthesized from 1 µg mRNA using a high-capacity cDNA reverse transcription kit (Applied Biosystems, #4368813). mRNA expression levels were measured with a 7500 Real-time PCR system (Applied Biosystems) using SYBR green dye (Life Technologies, #4367659). Primer sequences are listed in **Table S2**.

### *Ex vivo* efferocytosis assay

Primary mouse KCs were isolated, seeded in an 8-well chamber slide, and cultured at 37°C overnight. To induce apoptotic hepatocytes, mice were given Fas ligand (Jo2, BD Biosciences, # 554254, RRID:AB_395326, 100 μg/kg body weight) via intravenous injection for 2 hours. The hepatocytes were then isolated from the livers of these mice, cultured for 6 hours, and labeled with PKH67 (Sigma, #MINI67) according to the manufacturer’s protocol. PKH67-labeled apoptotic hepatocytes were incubated with KCs in the presence of anti-TIM4 antibody (20 μg/ml) or isotype IgG control for 6 hours. The KCs were then fixed in 4% paraformaldehyde, and F-actin was labeled with phalloidin-iFluor 594 reagent (Abcam, #ab176757). The images were captured using a Leica DMI 6000B fluorescence microscope, and the data were analyzed using Image J software. Primary human hepatocytes were treated with tumor necrosis factor-related apoptosis-inducing ligand (TRAIL, 200 ng/ml, Sigma, #T9701) for 6 hours to induce apoptosis. Apoptotic human hepatocytes were labeled with PKH67 and incubated with primary human KCs for 6 hours in the presence of anti-TIM4 or isotype control IgG. The KCs were then fixed in 4% paraformaldehyde, and F-actin was labeled with phalloidin-iFluor 594 reagent (Abcam, #ab176757). Images were captured using a green spinning-disk confocal microscope (Nikon Ti), and the data were analyzed using Imaris 9.5 quantification software.

### Statistical Analysis

All quantitative data are presented as mean ± SEM. Statistical significance was determined using GraphPad Prism software (version 9.3). The Shapiro-Wilk test was used to test normality. Statistical significance between the two groups was analyzed using the Student’s t-test, and multiple groups were analyzed using one-way ANOVA with Tukey post-hoc testing. P values less than 0.05 were considered statistically significant.

## Results

### Efferocytosis by liver macrophages is decreased in human and experimental NASH and is correlated with the loss of macrophage TIM4

Apoptosis in hepatocytes is a key feature of NASH, and accumulation of apoptotic hepatocytes has been reported in NASH liver^5,6^. As macrophage efferocytosis is crucial in clearing dead cells and maintaining tissue homeostasis, we postulated that the efferocytosis ability of liver macrophages is impaired in advanced NASH vs. early NASH. To evaluate our hypothesis, we utilized human liver biopsy specimens with various non-alcoholic fatty liver disease activity scores (NAS) and conducted an in situ efferocytosis assay, as described previously^29^. Based on measurements of the ratio of macrophage-associated:free TUNEL^+^ cells, efferocytosis by liver macrophages was markedly decreased in advanced NASH liver (NAS score 3-6) compared with early NASH liver (NAS score 0-2). In addition, the decrease in efferocytosis in NASH was accompanied by an increase in the total number of TUNEL^+^ cells (**Fig. 1A**). To further explore the potential mechanism underlying impaired efferocytosis in a mouse NASH model, we fed mice the high-fat choline-deficient L-amino acid-defined diet (HF-CDAA) diet for 2, 3, 4, 8, or 12 weeks. These mice develop hepatic steatosis at 2 weeks, early liver fibrosis at 4 weeks, and advanced liver fibrosis at 8 and 12 weeks of diet feeding^26^ (**Fig. S1A**). Based on the ratio of macrophage-associated:free cl-caspase 3^+^ (apoptotic) cells, efferocytosis by liver macrophages was markedly decreased at the 8-and 12-week time points, accompanied by an increase in cl-caspase 3^+^ cells (**Fig. 1B**).

One mechanism of impaired efferocytosis is loss of macrophage receptors that recognize apoptotic cells. We therefore measured mRNAs encoding efferocytosis receptors in 12 weeks HF-CDAA-fed mice vs. chow-fed mice. The mRNAs encoding MerTK, BAI1, and AXL were increased in NASH vs. control liver **(Fig. S1B, first 3 groups**), suggesting that the loss of these receptors was not involved in defective efferocytosis in NASH. The finding with *Mertk* is consistent with our previous demonstration that liver macrophage efferocytosis in mice fed the fructose-palmitate-cholesterol (FPC) diet for 16 weeks was not impaired by deletion of macrophage MerTK^30^, and we now show similar results using the HD-CDAA NASH model (**Fig. S1C**). However, consistent with previous reports in NASH mice^18-21^, the mRNA encoding the efferocytosis receptor TIM4 (*Timd4*) and TIM4 protein were decreased in the livers of both FPC- and HF-CDAA-induced NASH mice (**Fig. S1B, 4^th^ group, and Fig. S1C-I**). In particular, we observed a decrease in *Timd4* expression after 8 weeks of HF-CDAA feeding (**Fig. S1D**), which coincides with the point at which impaired liver macrophage efferocytosis occurs (above). In addition, both liver TIM4 protein by immunoblot and TIM4 expression in Clec4f^+^ KCs by immunofluorescence microscopy were decreased after 12 weeks of HF-CDAA feeding (**Fig. S1E,F**). Similar results were found in the livers of mice fed the FPC diet for 16 weeks (**Fig. S1G-I**). Consistent with the HF-CDAA time course data above, KC TIM4 expression was not decreased in 8-week FPC-fed mice (**Fig. S1J**), which have steatosis but not NASH^25^. Most importantly, as the expression TIM4 in human NASH had not been previously reported, we now show that TIM4 protein as measured by immunoblot was decreased in human NASH liver compared with normal liver (**Fig. 1C**). Further, immunofluorescence staining of human liver showed that the expression of TIM4 in liver CD68^+^ macrophages was decreased in NASH (**Fig. 1D**). In summary, mouse and human NASH show evidence of impaired efferocytosis, and this is associated with a decrease in TIM4 expression on liver macrophages. Based on these findings, we focused on the hypothesis that the decrease of macrophage TIM4 in NASH is a key mechanism for impaired efferocytosis and, consequently, progression to fibrotic NASH.

### Antibody-mediated TIM4 blockade and genetic deletion of KC TIM4 reduces efferocytosis and accelerates liver fibrosis in experimental NASH

To investigate the role of TIM4 in the clearance apoptotic hepatocytes by human KCs *ex vivo*, we incubated primary human KCs with fluorescently labeled Fas-induced apoptotic hepatocytes in the presence of anti-TIM4 *vs.* IgG control. Anti-TIM4 decreased efferocytosis as measured by either volume or mean fluorescence intensity (MFI) of engulfed apoptotic hepatocytes (**Fig. 2A**). Similar results were found using primary mouse liver KCs (**Fig. 2B**). To evaluate the role of TIM4 in early NASH, mice were fed a HF-CDAA diet for 4 weeks, with anti-TIM4 or IgG administered during weeks 3 and 4. The two groups of mice had similar body and liver weights (**Fig. S2A**). Anti-TIM4 administration decreased the ratio of macrophage-associated:free cl-caspase-3^+^ (apoptotic) cells, indicating impaired efferocytosis (**Fig. 2C**). Most importantly, anti-TIM4 increased liver fibrosis as assessed by both picrosirius red and collagen 1A1 (COL1A1) staining (**Fig. 2D,E**). Hepatic stellate cell activation was also increased by anti-TIM4 administration, as indicated by higher α-SMA^+^ and Opn^+^ areas (**Fig. 2G,H**). The livers of the anti-TIM4 cohort also showed increased CK19^+^ area, indicating the presence of bile ductular reaction (**Fig. 2H**), which is associated with NASH pathogenesis^31^ ^32^. Finally, anti-TIM4 administration increased the number of hepatic F4/80^+^ macrophage crown-like structures (hCLS) in the liver, which is a characteristic feature of mouse and human NASH associated with liver fibrosis^33^. In contrast, anti-TIM4 did not affect hepatosteatosis, plasma ALT activity, or liver macrophage as assessed by *Adgre1* mRNA level (**Fig. S2B-D**).

**Fig. 2.**
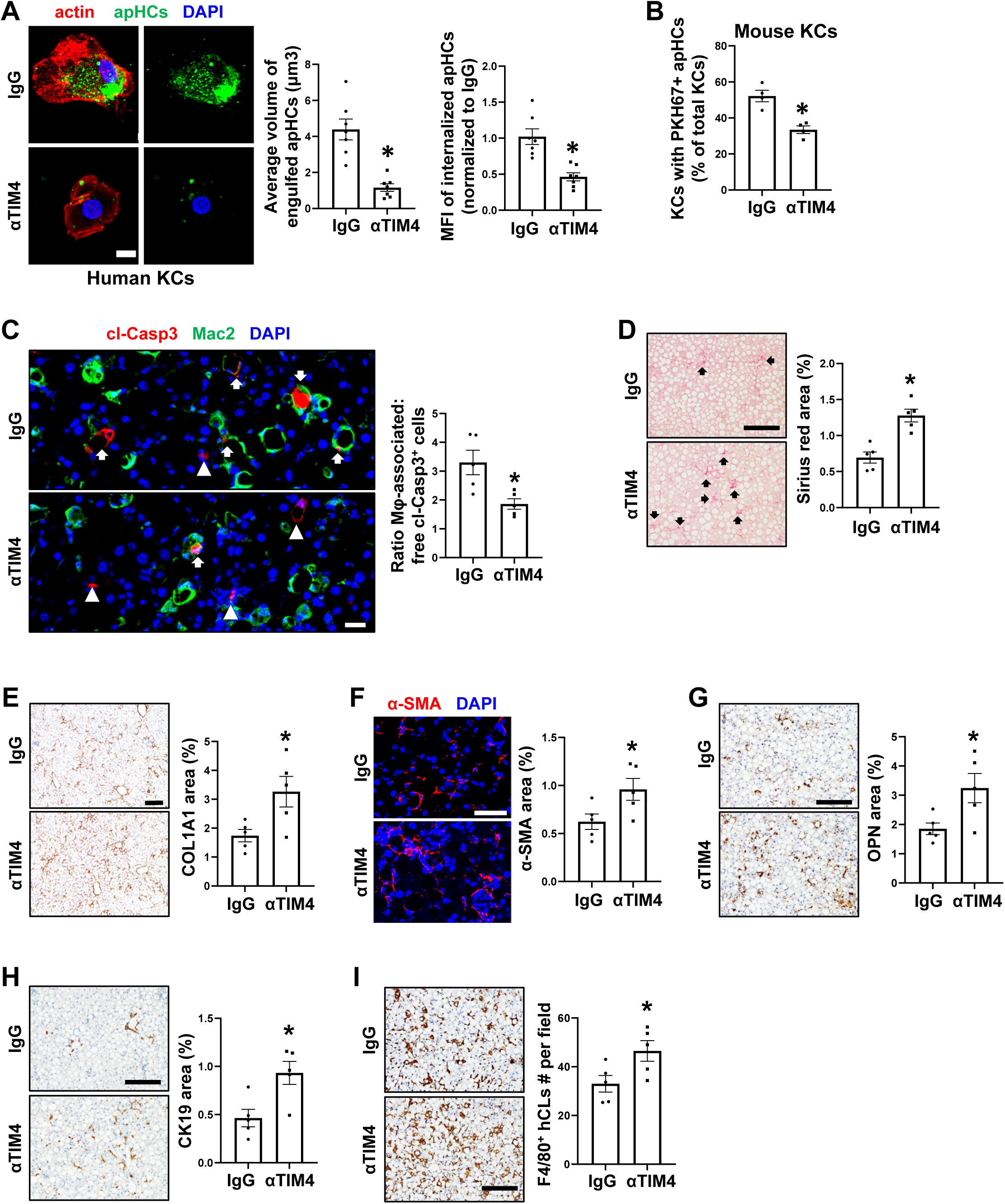
Anti-TIM4 reduces efferocytosis and accelerates liver fibrosis in early NASH. (**A**) Images of primary human KCs stained for F-actin (red) after exposure to apoptotic human hepatocytes (apHCs, green) for 6 h in the presence of IgG or anti-TIM4. Bar, 10 μm. The data were quantified as both the average volume of engulfed cargo and the MFI of internalized apHCs (n = 7 biological replicate cells/group). (**B**) Quantification of efferocytosis by primary mouse KCs after exposure to PKH67-labaled apoptotic mouse hepatocytes (n = 4 biological replicate cells/group). (**C-I**) Mice were fed the HF-CDAA NASH diet for 4 weeks and treated with IgG or anti-TIM4 during weeks 3-4 (n = 5 mice/group). (**C**) Images of liver sections immunostained with anti-Mac2 (green) and anti-cl-Casp3 (red). Bar, 25 μm. Arrows depict efferocytic macrophage, while arrowheads depict macrophage-free apoptotic cells. Efferocytosis was quantified as in Fig. 1B. (**D**) Staining and quantification of picrosirius red–positive area. Bar, 200 μm. (**E**) Immunostaining and quantification of collagen 1a1 (COL1A1)-positive area. Bar, 200 μm. (**F**) Immunostaining and quantification of α-SMA-positive area. Bar, 50 μm. (**G**) Immunostaining and quantification of osteopontin (OPN)-positive area. Bar, 200 μm. (**H**) Immunostaining and quantification of cytokeratin 19 (CK19)-positive area. Bar, 200 μm. (**I**) Immunostaining of F4/80, with quantification of F4/80^+^ hepatic crown-like structures (hCLS)/field. Bar, 200 μm. For immunofluorescence images, nuclei are stained with DAPI (blue). All data are means ± SEM. *p <0.05, **p <0.01 by Student’s t-test.

To investigate the role of KC TIM4 in liver efferocytosis and NASH progression, we generated *Timd4^fl/fl^* mice (**Fig. S2E**) and crossed them KC-specific with *Clec4f-Cre* mice^28^ to knockdown TIM4 in KCs (KC-TIM4-KD). *Timd4^fl/fl^* mice were used as the control mice. We first characterized chow-fed mice and found that liver *Timd4* mRNA expression was markedly decreased in KC-TIM4-KD mice, while the liver mRNAs of other efferocytosis receptors were similar between KD and control mice (**Fig. S2F**). Consistent with the *Timd4* data, immunofluorescence analysis showed markedly decreased TIM4 expression in liver macrophages in the KD cohort (**Fig. S2G**), while the expression of Clec4f was similar in control and KD liver (**Fig. S2H**). We then fed the mice the HF-CDAA diet for 4 weeks and verified decreased KC -TIM4 in the KD livers (**Fig. 3A**). The two cohorts had similar body and liver weights and fasting blood glucose (**Fig. S2I,J**), but KC-TIM4-KD was associated with defective efferocytosis by liver macrophages, indicated by a reduced ratio of macrophage-associated:free cl-caspase3^+^ cells and increased total cl-caspase3^+^ number per field (**Fig. 3B**). Most importantly, the KD mice had increased liver fibrosis based on picrosirius red and COL1A1 staining (**Fig. 3C,D**) and evidence of increased HSC activation based on higher α-SMA^+^ and Opn^+^ areas (**Fig. 3E,F**). As with the anti-TIM4 experiment, the KD mice had increased CK19+ area and hepatic crown-like structure (**Fig. 3G,H**). These changes occurred despite no change in hepatosteatosis and only a modest decrease in plasma ALT activity (**Fig. S2K,L**). KC-TIM4-KD also lowered efferocytosis and fibrosis in HF-CDAA-fed female mice without affect hepatosteatosis (**Fig. 3I-L and Fig. S3A-C**).

**Fig. 3.**
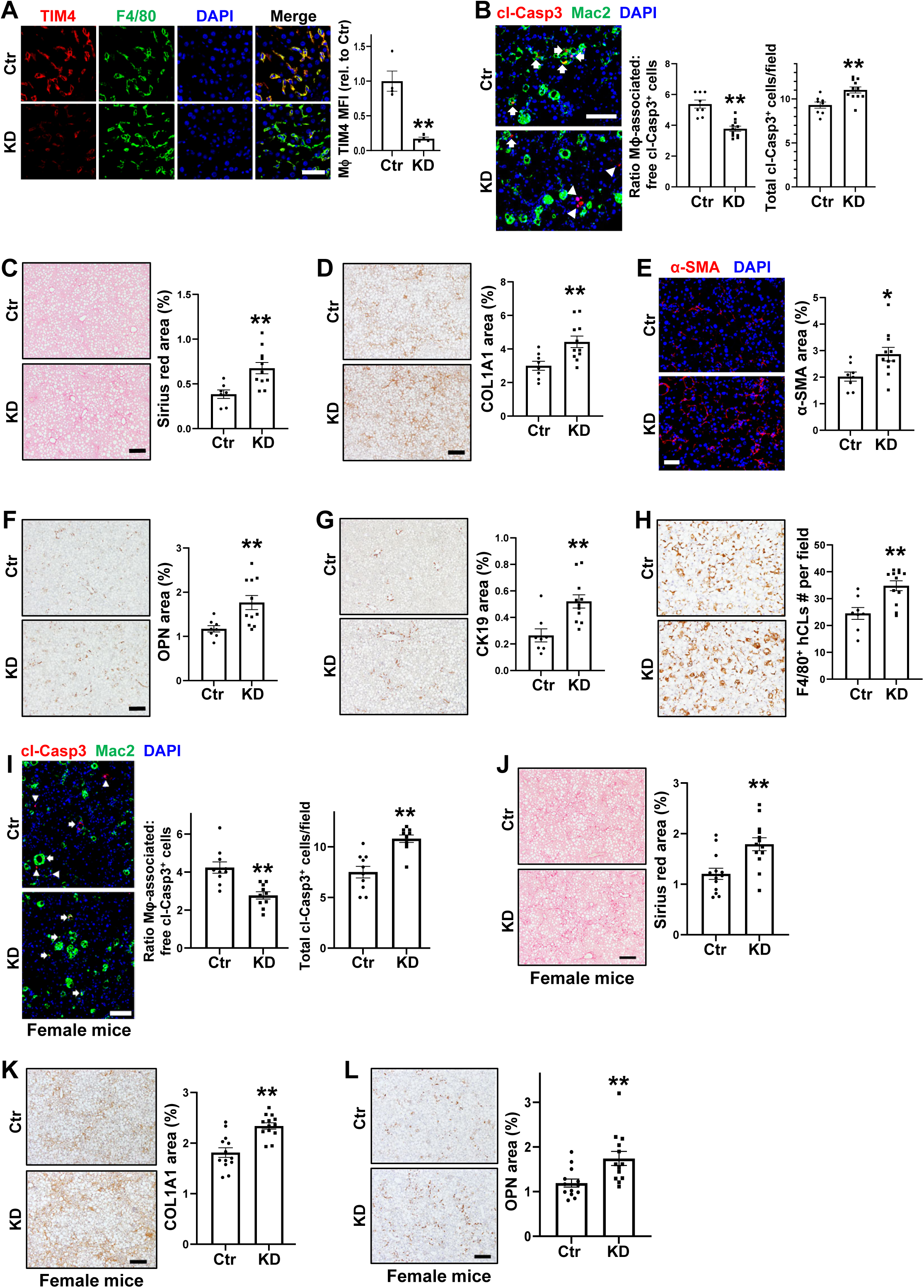
Knockdown of Kupffer cell TIM4 decreases efferocytosis and accelerates liver fibrosis in early HF-CDAA diet-induced NASH. For panels **A-H**, *Timd4^fl/fl^* (control, Ctr) and *Timd4^fl/fl^*;*Clec4f-Cre^+/-^* (knockdown, KD) male mice were fed the HF-CDAA diet for 4 weeks (n = 8-10 mice/group). (**A**) Immunostaining of liver sections using anti-TIM4 (red) and anti-F4/80 (green). Data were quantified as macrophage TIM4 MFI relative to control mice (n = 4 mice/group). Bar, 50 μm. (**B**) Images of liver sections immunostained with anti-Mac2 (green) and anti-cl-Casp3 (red). Bar, 50 μm. Arrows depict efferocytic macrophages, while arrowheads depict macrophage-free apoptotic cells. Efferocytosis was quantified as in Fig. 1B. (**C**) Staining and quantification of picrosirius red–positive area (arrows). Bar, 200 μm. (**D**) Immunostaining and quantification of collagen 1a1-positive area. Bar, 200 μm. (**E**) Immunostaining and quantification of α-SMA-positive area. Bar, 50 μm. (**F**) Immunostaining and quantification of OPN-positive area. Bar, 200 μm. (**G**) Immunostaining and quantification of CK19-positive area. Bar, 200 μm. (**H**) Immunostaining of F4/80, with quantification of F4/80^+^ hepatic crown-like structures/field. Bar, 200 μm. For panels **I-K**, *Timd4^fl/fl^*(control, Ctr) and *Timd4^fl/fl^*;*Clec4f-Cre^+/-^*(knockdown, KD) mice were fed the HF-CDAA diet for 8 weeks (n = 13 mice/group). (**I**) Images of liver sections immunostained with anti-Mac2 (green) and anti-cl-Casp3 (red). Bar, 100 μm. The arrows depicts an efferocytic macrophage, and the arrowheads depict macrophage-free apoptotic cells. Efferocytosis was quantified as in Fig. 1B. (**J**) Staining and quantification of picrosirius red–positive area (arrows). Bar, 200 μm. (**K**) Immunostaining and quantification of collagen 1a1-positive area. Bar, 200 μm. (**L**) Immunostaining and quantification of osteopontin (OPN)-positive area. Bar, 200 μm. For immunofluorescence images, nuclei are stained with DAPI (blue). All data are means ± SEM. *p <0.05, **p <0.01 by Student’s t-test.

To test the effect of KC-TIM4-KD in another NASH model, we fed KD and control mice the FPC diet for 16 weeks. As with the HF-CDAA model, we achieved robust KD of KC-TIM4 (**Fig. 4A**), and there were no significant differences between the two cohorts in body and liver weight, fasting blood glucose, hepatosteatosis, and plasma ALT (**Fig. S3D-G**). Most importantly, KC-TIM4-KD caused a decease in efferocytosis by liver macrophages (**Fig. 4B**) and increases in liver fibrosis (**Fig. 4C,D**), α-SMA^+^ and Opn^+^ areas (**Fig. 4E,F**), and mRNAs associated with HSC activation, *i.e.*, *Col1a1*, *Col1a2*, *Col3a1*, *Spp1*, and *Timp1* (**Fig. 4G**). The livers of the KD mice also had increases in CK19^+^ area (**Fig. 4H**) and hepatic crown-like structures (**Fig. 4I**). In contrast, mRNAs encoding proteins associated with inflammation were similar between the two cohorts (**Fig. S3H**). Thus, when KC-TIM4 is genetically lowered in early NASH, *i.e.*, before it is naturally lowered in later NASH, the progression to NASH fibrosis is accelerated.

**Fig. 4.**
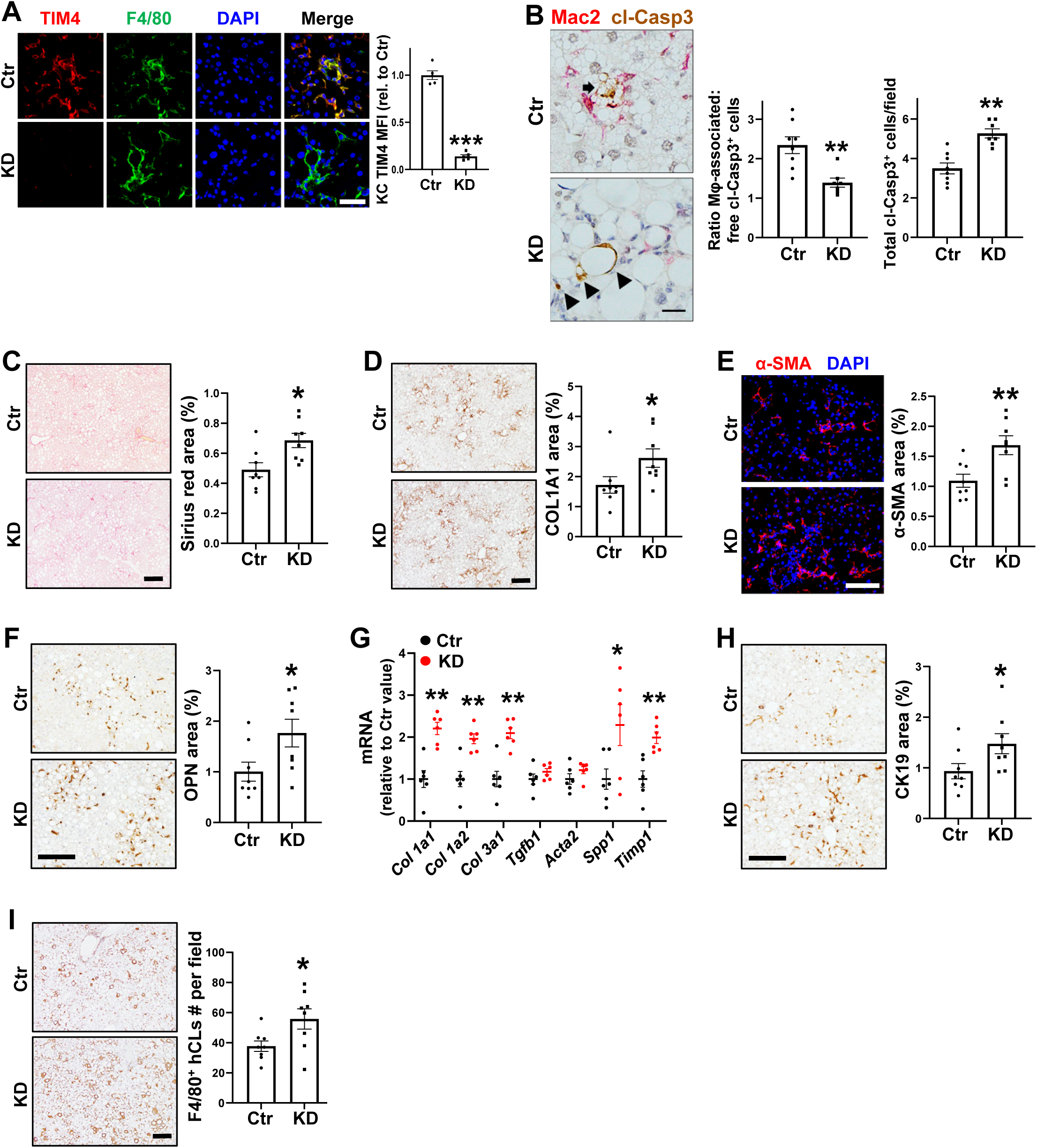
Knockdown of Kupffer cell TIM4 decreases efferocytosis and accelerates liver fibrosis in early FPC diet-induced NASH. *Timd4^fl/fl^* (control, Ctr) and *Timd4^fl/fl^*;*Clec4f-Cre* (knockdown, KD) male mice were fed the FPC diet for 16 weeks (n = 8-9 mice/group). (**A**) Immunostaining of liver sections using anti-TIM4 (red) and anti-F4/80 (green). Data were quantified as macrophage TIM4 MFI relative to control mice (n = 5 mice/group). Bar, 50 μm. (**B**) Images of liver sections immunostained with anti-Mac2 (green) and anti-cl-Casp3 (red). Bar, 50 μm. The arrow depicts an efferocytic macrophage, and the arrowheads arrows depict macrophage-free apoptotic cells. Efferocytosis was quantified as in Fig. 1B. (**C**) Staining and quantification of picrosirius red–positive area (arrows). Bar, 50 μm. (**D**) Immunostaining and quantification of COL1A1-positive area. Bar, 200 μm. (**E**) Immunostaining and quantification of α-SMA-positive area. Bar, 50 μm. (**F**) Immunostaining and quantification of OPN-positive area. Bar, 200 μm. (**G**) Liver mRNAs for the indicated fibrogenic genes, expressed as relative to the values for control mice. (**H**) Immunostaining and quantification of CK19-positive area. Bar, 200 μm. (**I**) Immunostaining of F4/80, with quantification of hepatic crown-like structures/field. Bar, 200 μm. All data are means ± SEM. *p <0.05, **p <0.01, ***p <0.001 by Student’s t-test.

### Macrophage TIM4 restoration after the development of steatosis promotes efferocytosis and suppresses the progression to liver fibrosis in NASH mice

The most important causation question related to TIM4, efferocytosis, and NASH is whether the *restoration* of macrophage TIM4 during the transition from early to advanced NASH would dampen the progression to liver fibrosis. To achieve this goal, we created a transgenic (Tg) model in which *Timd4* could be expressed in macrophages in a doxycycline (Dox)-inducible manner (iTg mice). We crossed mice transgenic for TRE (7 tetO+ minimal CMV promoter)-*Timd4*-bGHpA, with macrophage-specific *Cd68rtTA* mice, using mice with TRE-*Timd4*-bGHpa alone and *Cd68rtTA* as controls. The iTg and control mice were fed the HF-CDAA diet for 10 weeks, with Dox was administered in the drinking water between weeks 6 and 10. This protocol successfully restored TIM4 in liver macrophages (**Fig. 5A**). Restoration of macrophage TIM4 did not exert any influence on body weight, liver weight, and fasting blood glucose levels (**Fig. S4A,B**). Most importantly, the TIM4-restored NASH mice showed improved efferocytosis (**Fig. 5B**) and decreases in liver fibrosis (**Fig. 5C,D**), HSC activation (**Fig. 5E,F**), bile ductular reaction (**Fig. 5G**), and hepatic crown-like structures (**Fig. 5H**). These improvements occurred without any changes in hepatic steatosis or plasma ALT (**Fig. S4C,D**).

**Fig. 5.**
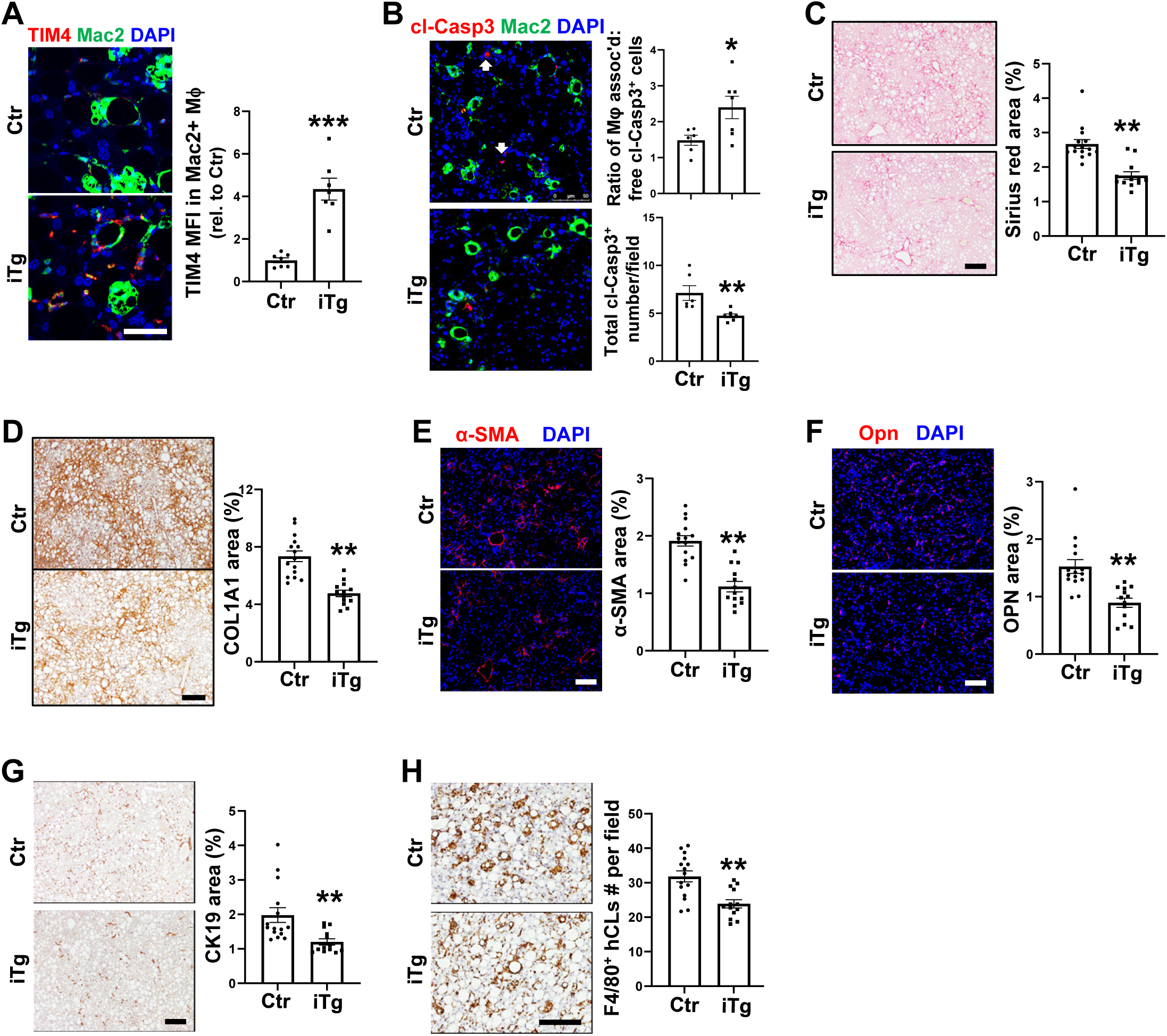
TIM4 restoration in liver macrophages in HF-CDAA diet-induced NASH promotes efferocytosis and lowers liver fibrosis. Control mice (TRE-TIMD4 and CD68rtTA; Ctr; n = 15) and inducible CD68rtTA:TRE-TIMD4 transgenic mice (iTg, n = 13) were fed the HF-CDAA diet for 10 weeks, with 75 μg/ml doxycycline (Dox) added to the drinking water during weeks 6-10. (**A**) Immunostaining of liver sections using anti-TIM4 (red) and anti-Mac2 (green). Data were quantified as macrophage TIM4 MFI relative to control mice. Bar, 50 μm. (**B**) Images of liver sections immunostained with anti-Mac2 (green) and anti-cl-Casp3 (red). Bar, 25 μm. Arrows depict macrophage-free apoptotic cells. Efferocytosis was quantified as in Fig. 1B. (**C**) Staining and quantification of picrosirius red– positive area (arrows). Bar, 200 μm. (**D**) Immunostaining and quantification of COL1A1-positive area. Bar, 200 μm. (**E**) Immunostaining and quantification of α-SMA-positive area. Bar, 100 μm. (**F**) Immunostaining and quantification of OPN-positive area. Bar, 100 μm. (**G**) Immunostaining and quantification of CK19-positive area. Bar, 200 μm. (**H**) Immunostaining of F4/80, with quantification of hepatic crown-like structures/field. Bar, 200 μm. All data are means ± SEM. *p <0.05, **p <0.01, ***p <0.001 by Student’s t-test.

We then conducted a similar experiment in 16-week FPC-fed mice, intervening with Dox between weeks 9-16 so that macrophage TIM4 would be restored during steatosis-to-NASH progression, which is the period during TIM4 expression and efferocytosis decrease (above) and liver fibrosis increases^25^. The results were similar to those with the HF-CDAA model. There was clear restoration of liver macrophage TIM4 in the Dox-treated cohort (**Fig. 6A**), and this led to improvements in liver macrophage efferocytosis (**Fig. 6B**) and decreases in liver fibrosis (**Fig. 6C,D**), α-SMA^+^ and OPN^+^ areas (**Fig. 6E,F**), mRNAs associated with HSC activation (*Col1a1*, *Col1a2*, *Col1a3*, *Tgfb1*, and *Timp1*) (**Fig. 6G**), CK19^+^ areas (**Fig. 6H**), and hepatic crown-like structures (**Fig. 6I**)—all without changes in body weight, liver weight, fasting blood glucose, hepatic steatosis, or plasma ALT (**Fig. S4E-H**). In contrast, mRNAs encoding proteins associated with inflammation were similar between the two cohorts (**Fig. S4I**). The findings, within the context of the previous data in this report, provide strong evidence that the decrease in TIM4 in liver macrophages during NASH progression contributes to impaired efferocytosis and the progression of liver fibrosis.

**Fig. 6.**
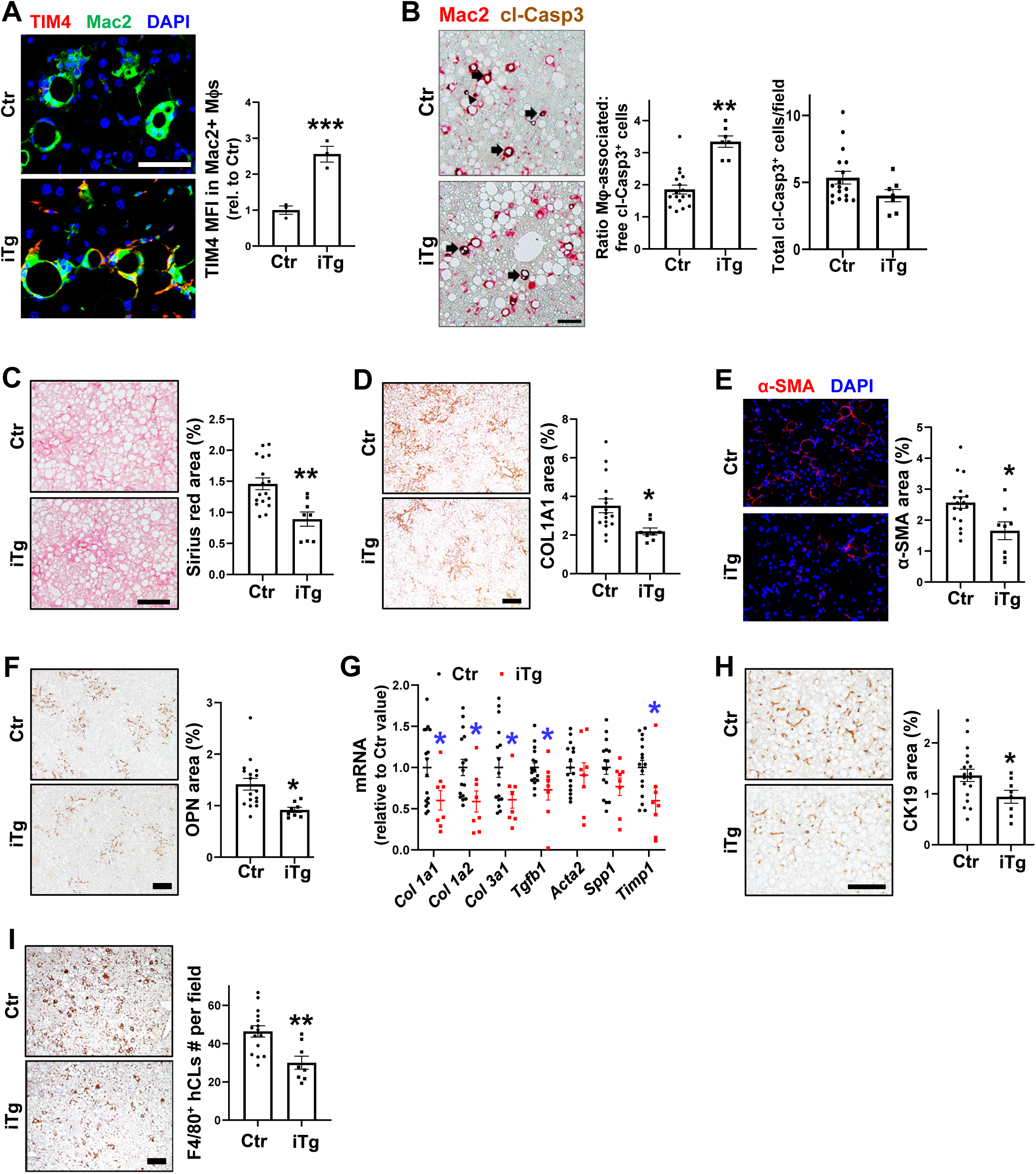
TIM4 restoration in liver macrophages in FPC diet-induced NASH promotes efferocytosis and lowers liver fibrosis. (**A-H**) Control mice (TRE-TIMD4 and CD68rtTA; Ctr; n = 17) and inducible CD68rtTA:TRE-TIMD4 transgenic mice (iTg, n = 8) were fed the FPC diet for 16 weeks, with 75 μg/ml doxycycline (Dox) added to the drinking water during weeks 9-16. (**A**) Immunostaining of liver sections using anti-TIM4 (red) and anti-Mac2 (green). Data were quantified as macrophage TIM4 MFI relative to control mice (n = 3 mice/group). Bar, 50 μm. (**B**) Images of liver sections immunostained with anti-Mac2 (green) and anti-cl-Casp3 (red). Bar, 100 μm. Arrows depict efferocytic macrophages, and the arrowhead depicts a macrophage-free apoptotic cell. Efferocytosis was quantified as in Fig. 1B. (**C**) Staining and quantification of picrosirius red–positive area (arrows). Bar, 100 μm. (**D**) Immunostaining and quantification of COL1A1-positive area. Bar, 200 μm. (**E**) Immunostaining and quantification of α-SMA-positive area. Bar, 50 μm. (**F**) Immunostaining and quantification of OPN-positive area. Bar, 200 μm. (**G**) Liver mRNAs for the indicated fibrogenic genes, expressed as relative to the values for control mice. (**H**) Immunostaining and quantification of CK19-positive area. Bar, 200 μm. (**I**) Immunostaining of F4/80, with quantification of F4/80^+^ crown-like structures/field. Bar, 200 μm. All data are means ± SEM. *p <0.05, **p <0.01, ***p <0.001 by Student’s t-test.

## Discussion

A key feature of NASH is hepatocyte apoptosis, which correlates with the clinically most important feature of the disease, liver fibrosis^5,6^. Therefore, a key area in NASH research is to understand the fate of apoptotic hepatocytes and how hepatocyte apoptosis might be linked to liver fibrogenesis. Addressing these fundamental issues may suggest novel strategies to prevent fibrosis progression in NASH. In terms of apoptotic hepatocyte fate, the finding that they increase in advance human and experimental NASH indicates that there is a defect in their clearance, as apoptotic cells in tissues do not accumulate if efferocytosis is intact unless massive apoptosis occurs, which is not the case in NASH. Although a recent report showed that apoptotic hepatocytes accumulate in liver when another macrophage efferocytosis receptor was deleted^12^, there has been no previous report directly measuring efferocytosis in liver, *i.e.*, macrophage-associated *vs*. free apoptotic cells, during NASH progression in mice or humans. The previous findings that TIM4 expression is decreased in mouse NASH are consistent with its possible protective role, and our new finding that liver macrophage TIM4 is decreased and correlates with defective efferocytosis in human NASH bolsters the central concept herein. Moreover, TIM4 was shown to mediate apoptotic hepatocyte uptake by KCs *ex vivo*^22-24^, but its role in efferocytosis by liver macrophages and liver fibrosis in NASH has remained unknown. There has been recent interest in TREM2, which is expressed on liver macrophages and can mediate efferocytosis *ex vivo*^12^. However, although there is some evidence that TREM2 may mediate efferocytosis in experimental NASH, the expression levels and roles of TREM2 in efferocytosis and liver fibrosis in NASH remain controversial^12-15^. Finally, we showed previously that the clearance of necroptotic hepatocytes is defective in NASH owing to increases in SIRPα on NASH liver macrophages and CD47 on necroptotic heptocytes^16^. While blocking either SIRPα or CD47 improved necroptotic hepatocyte uptake and dampened NASH progression, these manipulations did not lower apoptotic heptocytes^16^. We therefore conclude that defective clearance of both necroptotic and apoptotic hepatocytes, which is caused by distinct mechanisms, both contribute to NASH progression.

The major mechanistic question arising from the new findings herein is why defective clearance of apoptotic hepatocytes in NASH promotes liver fibrosis. Interestingly, the effect was specific for fibrosis, as steatosis, inflammation, and plasma ALT were not affected in any of the models. In theory, the accumulation of apoptotic hepatocytes could directly contribute to fibrosis, *e.g.*, via their release of fibrogenic factors^7,8^. However, the failure of apoptosis inhibitors to block NASH fibrosis in humans^9^ suggests the possibility of alternative mechanisms. For example, efferocytosis re-programs macrophages to promote tissue resolution^10,11^, and it is possible that efferocytosis-mediated re-programming promotes liver macrophage secretion of pro-fibrogenic factors and/or dampens the secretion of anti-fibrogenic factors. Of note, two known efferocytosis-induced mediators are interleukin-10^34^ and insulin-like growth factor-1^35^, both of which can inhibit HSC activation and reduce fibrogenesis^36-38^. As efferocytosis in other settings can *promote* fibrosis as part of the healing process, *e.g.*, via transforming growth factor-β (TGFβ)^10,11^, efferocytosis-induced macrophage re-programming may be distinct in the unique setting of NASH^39^. Indeed, we showed that restoration of TIM4-mediated efferocytosis in NASH decreased liver *Tgfb1* (Fig. 6I). Future studies will be needed to further investigate this issue.

The new insights gained from this study could have translational impact^40^. The decreased expression of TIM4 in advanced NASH may be attributed to the loss of *Timd4* expression in KCs and/or the loss of KCs themselves^18-21^. A previous study showed that infusion of KCs mitigates liver fibrosis in a mouse liver injury model. It may also be possible to genetically restore macrophage TIM4, e.g., by using macrophage-targeted nanoparticles containing *Tim4* mRNA. Another approach might lie in pharmacologically preventing TIM4 down-regulation in NASH. Although the mechanism in NASH is not known, P38 mitogen-activated protein kinase activation mediates the age-related decrease in *Timd4* expression in blister monocytes/macrophages, resulting in defective efferocytosis and inflammation resolution^41^. Moreover, deleting myeloid P38α or P38γ/δ improves diet induced NAFLD/NASH^42,43^. Although the mechanism of this effect was not linked to TIM4 or efferocytosis, it is possible that liver macrophage-targeted inhibition or silencing of P38 or a key downstream mediator could prevent TIM4 down-regulation and thereby improve efferocytosis and decrease liver fibrosis in NASH. Additionally, understanding the epigenetic signals that regulate *Timd4* expression may afford other opportunities for therapeutic intevention^44^. More generally, the evidence herein that macrophage TIM4 down-regulation in NASH is not simply a biomarker of NASH macrophages but rather actively promotes liver fibrosis provides the impetus for future studies on the mechanism of *Timd4* regulation and how it may be amenable to therapeutic intervention.

## Abbreviations

ALT: alanine transaminase
HF-CDAA: high-fat, choline-deficient, L- amino acid-defined
HSC: hepatic stellate cell
FPC: fructose-palmitate-cholesterol
NAFLD: nonalcoholic fatty liver disease
KC: Kupffer cell
NASH: non-alcoholic steatohepatitis
TIM4: T cell immunoglobulin and mucin domain containing 4
TUNEL: terminal deoxynucleotidyl transferase dUTP nick end labeling

## Author contributions

H.S., X.W., B.G., A.Y., Jr., M.P.M., and I.T. participated in the conception and experimental design of the study. H.S. conducted the mouse experiments and liver analyses. L.V., A.J.S, and F.M. contributed to the human normal and NASH liver studies. C.S.L designed and generated the Timd4 knockdown and inducible transgenic models. H.S. and I.T. drafted the manuscript. All authors revised the manuscript and approved the final version.

## Data availability statement

Data are available from the corresponding authors upon reasonable request.

## Acknowledgements

We thank Drs. David Ngai and Maaike Schilperoort (CUIMC) for manuscript editing and Dr. Robert Schwabe (CUIMC) for helpful discussions.

