## Supplemental files and data for "Loss of TIM4-Dependent Efferocytosis in Kupffer Cells Promotes Liver Fibrosis in Nonalcoholic Steatohepatitis"

Fig. S1

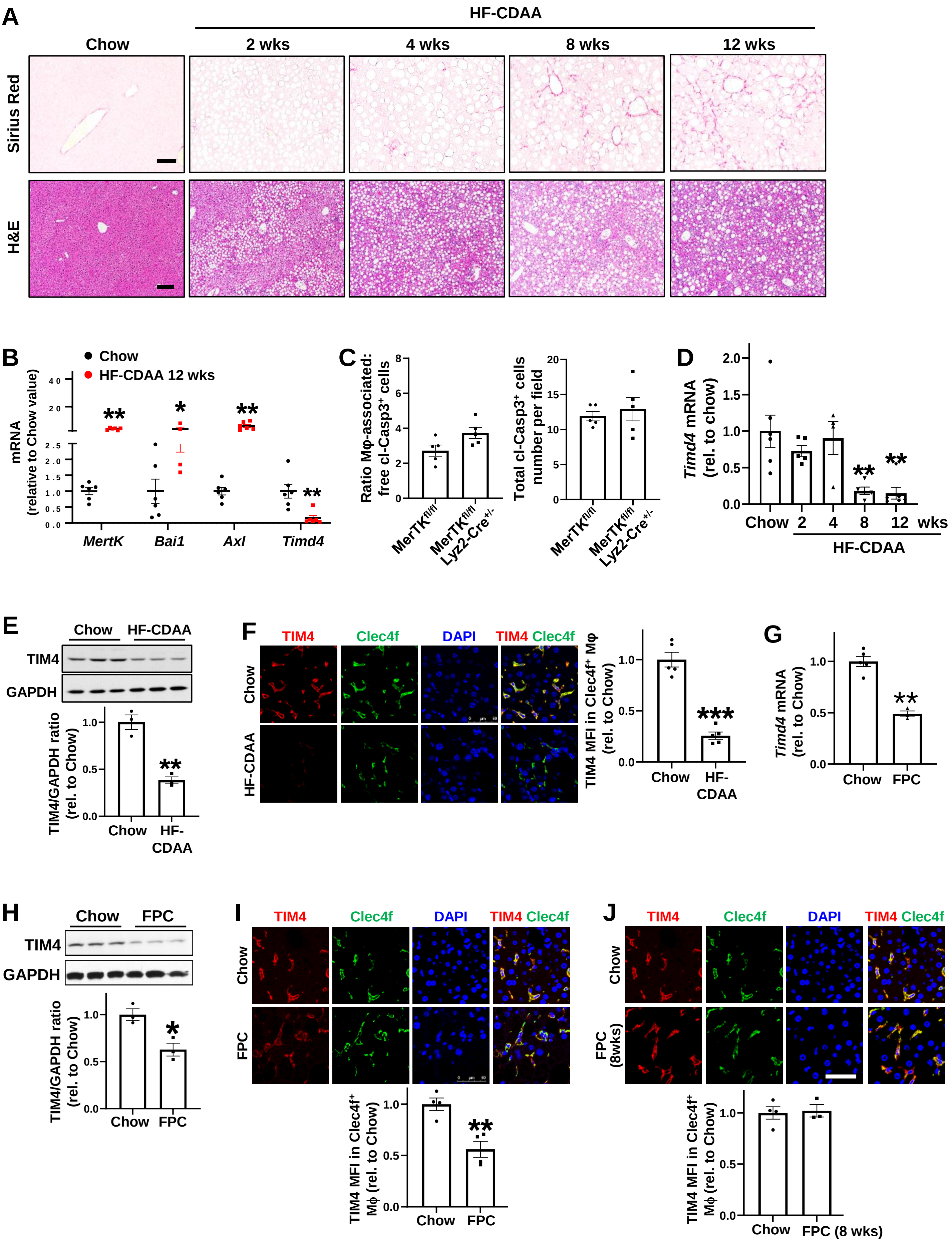

A
HF-CDAA
Chow
2 wks
4 wks
8 wks
12 wks
Sirius Red
H&E
B
Chow
HF-CDAA 12 wks
mRNA (relative to Chow value)
MertK
Bai1
Axl
Timd4
C
Ratio Mφ-associated: free cl-Casp3+ cells
Total cl-Casp3+ cells number per field
MerTKfl/fl
MerTKfl/fl Lyz2-Cre+/-
D
Timd4 mRNA (rel. to chow)
Chow
2
4
8
12
wks
HF-CDAA
E
Chow
HF-CDAA
TIM4
GAPDH
TIM4/GAPDH ratio (rel. to Chow)
F
TIM4
Clec4f
DAPI
TIM4 Clec4f
Chow
HF-CDAA
TIM4 MFI in Clec4f+ Mφ (rel. to Chow)
G
Timd4 mRNA (rel. to Chow)
Chow
FPC
H
Chow
FPC
TIM4
GAPDH
TIM4/GAPDH ratio (rel. to Chow)
I
TIM4
Clec4f
DAPI
TIM4 Clec4f
Chow
FPC
TIM4 MFI in Clec4f+ Mφ (rel. to Chow)
J
TIM4
Clec4f
DAPI
TIM4 Clec4f
Chow
FPC (8wks)
TIM4 MFI in Clec4f+ Mφ (rel. to Chow)
Chow
FPC (8 wks)

**Fig. S1. Additional data documenting decreased liver macrophage TIM4 and efferocytosis in mouse NASH liver. Related to Fig. 1.** (**A**) Mice were fed chow diet or the HF-CDAA diet for the indicated times, and liver tissues were stained with picrosirius red and H&E. Bar, 50 µm for picrosirius red and 200 µm for H&E staining. (**B**) The livers of mice fed chow diet or the HF-CDAA diet for 12 weeks were assayed for mRNAs encoding the indicated efferocytosis receptors (n = 6 mice/group). (**C**) Livers from *Mertk*$^{fl/fl}$ and *Mertk*$^{fl/fl}$*;Lyz2-Cre*$^{+/-}$ mice fed the HF-CDAA diet for 4 weeks were assayed for efferocytosis as in Fig. 1B (n = 5 mice/group). (**D**) The livers of mice fed chow diet or the HF-CDAA diet for the indicated number of weeks were assayed for *Timd4* mRNA and expressed relative to the chow value (n = 4-6 mice/group). (**E**) Immunoblot of TIM4 in the livers of mice fed chow diet or the HF-CDAA diet for 12 weeks, with data quantification expressed relative to the chow value (n =3 mice/group). (**F**) Immunostaining of liver sections using anti-TIM4 (red) and anti-Clec4f (Kupffer cells, green) in the livers of mice fed chow diet or the HF-CDAA diet for 12 weeks, with the data expressed relative to the chow value (n = 5 mice per group). Bar, 50 µm. (**G-H**) *Timd4* mRNA and TIM4 protein in the livers of mice fed chow diet or the FPC diet for 16 weeks, expressed relative to the chow value (n = 3-5 mice/group). (**I**) The liver sections from mice fed chow diet or the FPC diet for 16 weeks were immunostained with anti-TIM4 (red) and anti-Clec4f (green), with data quantification expressed relative to the chow value (n = 4 mice/group). Bar, 50 µm. (**J**) The liver sections from mice fed chow diet or the FPC diet for 8 weeks were immunostained with anti-TIM4 (red) and anti-Clec4f (green), with data quantification expressed relative to the chow value (n = 3-4 mice/group). Bar, 50 µm. All data are means ± SEM. *p <0.05, **p <0.01, ***p <0.001 by Student's t-test for all panels except panel D, which used one-way ANOVA.

### Fig. S2

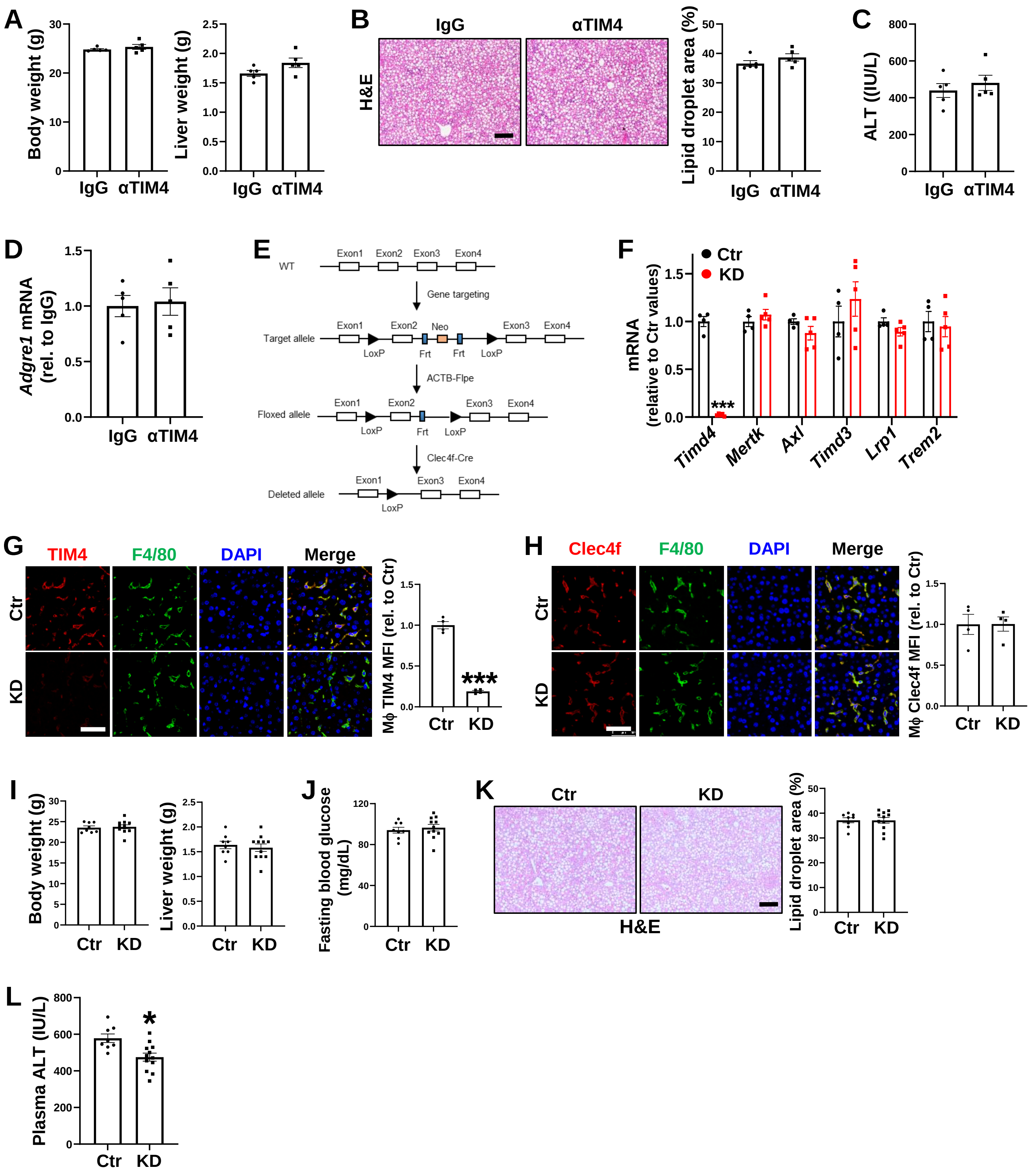

A
Body weight (g)
Liver weight (g)
IgG αTIM4
B
IgG
αTIM4
H&E
Lipid droplet area (%)
C
ALT ((IU/L)
D
Adgre1 mRNA (rel. to IgG)
E
WT
Exon1 Exon2 Exon3 Exon4
Gene targeting
Target allele
Neo
LoxP Frt Frt LoxP
ACTB-Flpe
Floxed allele
Clec4f-Cre
Deleted allele
F
Ctr
KD
mRNA (relative to Ctr values)
Timd4 Mertk Axl Timd3 Lrp1 Trem2
G
TIM4 F4/80 DAPI Merge
Ctr
KD
Mϕ TIM4 MFI (rel. to Ctr)
H
Clec4f F4/80 DAPI Merge
Mϕ Clec4f MFI (rel. to Ctr)
I
Body weight (g)
Liver weight (g)
J
Fasting blood glucose (mg/dL)
K
Ctr KD
H&E
Lipid droplet area (%)
L
Plasma ALT (IU/L)

**Fig. S2. Additional data documenting that blocking or genetic targeting of TIM4 decreases liver macrophage efferocytosis and accelerates liver fibrosis in early NASH. Related to Figs. 2-3.** (**A-D**) The IgG- and anti-TIM4-treated HF-CDAA early NASH mice described in Fig. 2 (n = 5 mice/group) were assayed for the following: (**A**) Body and liver weight; (**B**) H&E staining of liver sections and lipid droplet area quantification. Bar, 250 µm; (**C**) Plasma ALT activity; and (**D**) Liver *Adgre1* mRNA, expressed relative to the IgG value. (**E**) Strategy for generation of $Timd4^{fl/fl}$ mice. (**F-H**) The livers of 8-week chow-fed $Timd4^{fl/fl}$ (control, Ctr) and $Timd4^{fl/fl}$;*Clec4f-Cre*$^{+/-}$ (knockdown, KD) mice were assayed as follows (n=4-5 mice/group): (**F**) mRNAs encoding the indicated efferocytosis receptors; (**G**) TIM4 MFI in macrophages (Mϕs); and (**H**) Clec4f MFI in macrophages. All data are expressed relative to the control values. Bars, 50 µm. (**I-L**) HF-CDAA diet-fed control and TIM4-KD male mice described in Fig. 3A-H (n = 8-11 mice/group) were assayed for the following: (**I**) Body and liver weight; (**J**) Fasting blood glucose; (**K**) H&E staining of liver sections and lipid droplet area quantification. Bar, 250 µm; and (**L**) Plasma ALT activity. All data are means ± SEM. *p <0.05 by Student's t-test.

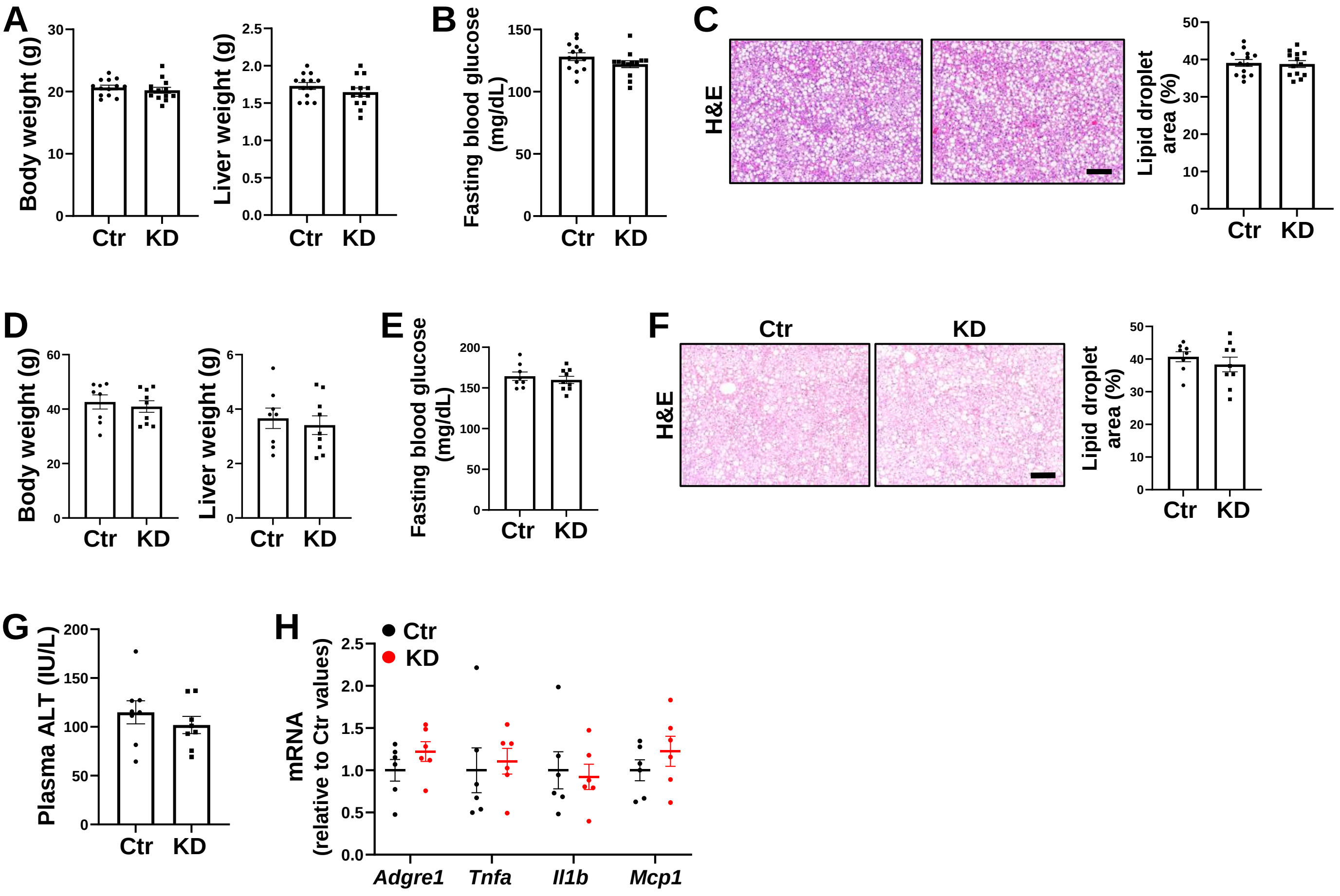
A
Body weight (g)
Liver weight (g)
B
Fasting blood glucose (mg/dL)
C
H&E
Lipid droplet area (%)
D
Body weight (g)
Liver weight (g)
E
Fasting blood glucose (mg/dL)
F
Ctr
KD
H&E
Lipid droplet area (%)
G
Plasma ALT (IU/L)
H
Ctr
KD
mRNA (relative to Ctr values)
Adgre1
Tnfa
Il1b
Mcp1

**Fig. S3. Additional data documenting that genetic targeting of TIM4 decreases liver macrophage efferocytosis and accelerates liver fibrosis in early NASH. Related to Figs. 3-4.** (**A-C**) HF-CDAA diet-fed control (Ctr) and TIM4-KD female mice described in Fig. 3I-K (n = 13 mice/group) were assayed for the following: (**A**) Body and liver weight; and (**B**) Fasting blood glucose. (**C**) H&E staining of liver sections and lipid droplet area quantification. Bar, 200 µm. (**D-H**) FPC diet-fed control (Ctr) and TIM4-KD mice described in Fig. 4 (n = 8-9 mice/group) were assayed for the following: (**D**) Body and liver weight; (**E**) Fasting blood glucose; (**F**) H&E staining of liver sections and lipid droplet area quantification. Bar, 250 µm; and (**G**) Plasma ALT activity. (**H**) Liver mRNAs encoding the indicated inflammation-related genes, expressed relative to the control values. All data are means ± SEM.

Fig. S4

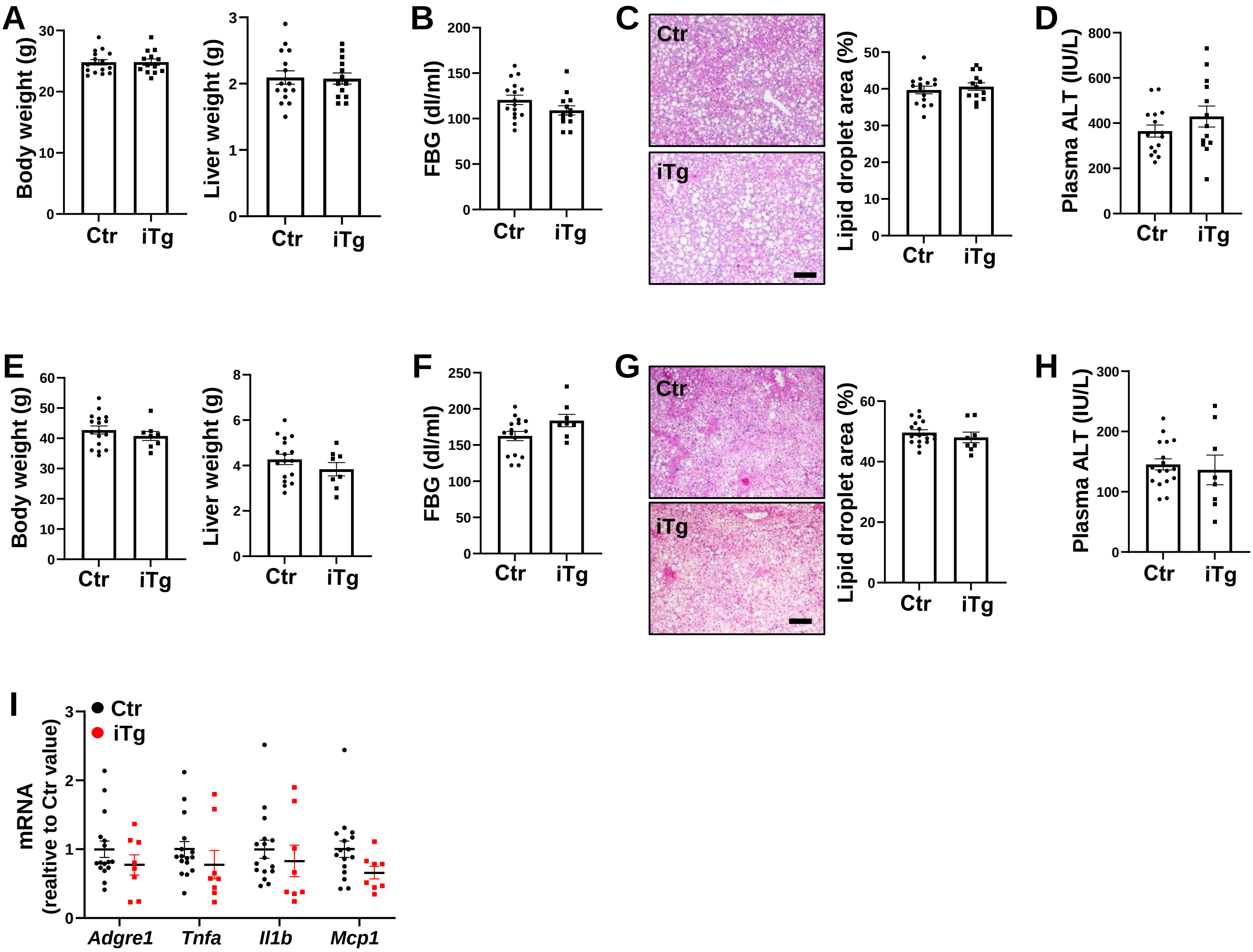
A
Body weight (g)
Ctr
iTg
Liver weight (g)
B
FBG (dl/ml)
C
Ctr
iTg
Lipid droplet area (%)
D
Plasma ALT (IU/L)
E
Body weight (g)
Liver weight (g)
F
FBG (dl/ml)
G
Lipid droplet area (%)
H
Plasma ALT (IU/L)
I
Ctr
iTg
mRNA
(realtive to Ctr value)
Adgre1
Tnfa
Il1b
Mcp1

**Fig. S4. Additional data documenting that macrophage TIM4 restoration promotes efferocytosis and lowers liver fibrosis in experimental NASH, Related to Figs. 5-6.** (**A-D**) The HF-CDAA-fed control and inducible TIM4 transgenic mice described in Fig. 5 (n = 13-15 mice/group) were assayed for the following: (**A**) Body and liver weight; (**B**) Fasting blood glucose; (**C**) H&E staining of liver sections and lipid droplet area quantification. Bar, 250 µm; and (**D**) Plasma ALT activity. (**E-H**) The FPC-fed control and inducible TIM4 transgenic mice described in Fig. 6 (n = 8-9 mice/group) were assayed for the following: (**E**) Body and liver weight; (**F**) Fasting blood glucose; (**G**) H&E staining of liver sections and lipid droplet area quantification. Bar, 250 µm; (**H**) Plasma ALT activity; and (**I**) Liver mRNAs encoding the indicated inflammation-related genes, expressed relative to the control values. All data are means ± SEM.

**Table S1 (related to Figure 1A, 1C, and 1D). Human liver samples.**

| ID | Age | Gender | Pathological diagnoses | Source (Figure) |
|---|---|---|---|---|
| HH532 | 48 | Male | Normal liver | Liver Tissue Cell Distribution System (Fig. 1C,D) |
| HH577 | 58 | Female | Normal liver | |
| HH698 | 46 | Female | Normal liver | |
| HH976 | 50 | Female | Normal liver | |
| HH996 | 55 | Male | Normal liver | |
| HH1020 | 45 | Male | Normal liver | |
| HH1069 | 41 | Female | Normal liver | |
| UMN1673 | 57 | Female | NASH cirrhosis | |
| UMN1253 | 65 | Female | NASH-no longer fatty. Cirrhosis, mildly active, with extensive cholestasis. Occasional Mallory bodies. | |
| UMN1259 | 56 | Male | NASH-almost no fat. End stage cirrhotic liver disease. Negative for malignancy. | |
| UMN1422 | 46 | Female | NASH-no longer fatty. Moderately active cirrhosis. No evidence of malignancy. Minimal current steatosis. | |
| UMN1443 | 68 | Male | NASH; cirrhosis, mixed macronodular and micronodular type. Minimal inflammatory activity, with minimal or no ongoing steatosis. No evidence of hepatocyte dysplasia or malignancy | |
| UMN1615 | 56 | Female | Advanced chronic liver disease (stage 4: "cirrhosis"). Marked fibrosis throughout the entire liver. The residual hepatocellular parenchyma shows abundant well-formed Mallory hyaline, consistent with advanced chronic liver disease from NASH. | |
| UMN1509 | 68 | Male | Cirrhosis with chronic hepatitis, mildly active, consistent with clinical history of NASH. Minimal steatosis, macrovesicular type (<5%). Viable resection margins. | |
| 19I-8031 | 69 | Male | NAS score = 5; Fibrosis score = 3 | Fondazione IRCCS Ca' Granda Ospedale Policlinico (Fig. 1A) |
| 17I-12781 | 56 | Male | NAS score = 3; Fibrosis score = 1c | |
| 17I-10585 | 53 | Female | NAS score = 3; Fibrosis score = 1c | |
| 18I-5368 | 51 | Female | NAS score = 3; Fibrosis score = 0 | |
| 19I-991 | 64 | Male | NAS score = 5; Fibrosis score = 2 | |
| 1n | 38 | Female | NAS score = 6; Fibrosis score = 1 | Virginia Common-wealth University (Fig. 1A) |
| 5n | 55 | Male | NAS score = 4; Fibrosis score = 2 | |
| 8n | 59 | Female | NAS score = 5; Fibrosis score = 3 | |
| 9n | 59 | Female | NAS score = 4; Fibrosis score = 3 | |
| 10n | 53 | Female | NAS score = 4; Fibrosis score = 2 | |
| 13n | 53 | Female | NAS score = 5; Fibrosis score = 2 | |
| 14n | 56 | Female | NAS score = 5; Fibrosis score = 1 | |
| 15n | 68 | Female | NAS score = 6; Fibrosis score = 3 | |
| 26C | 34 | Female | NAS score = 0; Fibrosis score = 0 | |
| 28C | 48 | Male | NAS score = 0; Fibrosis score = 1 | |
| 29C | 46 | Female | NAS score = 0; Fibrosis score = 0 | |
| 30C | 63 | Male | NAS score = 2; Fibrosis score = 0 | |

**Table S2.** Mouse primer sequence used for qPCR.

| Name | Sequence |
|---|---|
| *Hprt* F | TCAGTCAACGGGGGACATAAA |
| *Hprt* R | GGGGCTGTACTGCTTAACCAG |
| *Col1a1* F | GCTCCTCTTAGGGGCCACT |
| *Col1a1* R | CCACGTCTCACCATTGGGG |
| *Col1a2* F | GTAACTTCGTGCCTAGCAACA |
| *Col1a2* R | CCTTTGTCAGAATACTGAGCAGC |
| *Col3a1* F | CTGTAACATGGAAACTGGGGAAA |
| *Col3a1* R | CCATAGCTGAACTGAAAACCACC |
| *Tgfb1* F | CTCCCGTGGCTTCTAGTGC |
| *Tgfb1* R | GCCTTAGTTTGGACAGGATCTG |
| *Acta2* F | ATGCTCCCAGGGCTGTTTTCCCAT |
| *Acta2* R | GTGGTGCCAGATCTTTTCCATGTCG |
| *Spp1* F | CTGACCCATCTCAGAAGCAGAATCT |
| *Spp1* R | TCCATGTGGTCATGGCTTTCATTGG |
| *Timp1* F | GCAACTCGGACCTGGTCATAA |
| *Timp1* R | CGGCCCGTGATGAGAAACT |
| *Adgre1* F | ACCACAATACCTACATGCACC |
| *Adgre1* R | AAGCAGGCGAGGAAAAGATAG |
| *Tnfa* F | CTTCTGTCTACTGAACTTCGGG |
| *Tnfa* R | CAGGCTTGTCACTCGAATTTTG |
| *Il1b* F | CAACCAACAAGTGATATTCTCCATG |
| *Il1b* R | GATCCACACTCTCCAGCTGCA |
| *Mcp1* F | TTAAAAACCTGGATCGGAACCAA |
| *Mcp1* R | GCATTAGCTTCAGATTTACGGGT |
| *Timd3 F* | TCAGGTCTTACCCTCAACTGTG |
| *Timd3 R* | GGGCAGATAGGCATTTTTACCA |
| *Timd4 F* | CCTTCACTACAGAATCAGAAACTC |
| *Timd4 R* | CGGCTATGTCTGTAGATATGGT |
| *MerTK F* | CAGGGCCTTTACCAGGGAGA |
| *MerTK R* | TGTGTGCTGGATGTGATCTTC |
| *Axl F* | ATGGCCGACATTGCCAGTG |
| *Axl R* | CGGTAGTAATCCCCGTTGTAGA |
| *Lrp1 F* | ACTATGGATGCCCCTAAAACTTG |
| *Lrp1 R* | GCAATCTCTTTCACCGTCACA |
| *Trem2 F* | ACAGCACCTCCAGGAATCAAG |
| *Trem2 R* | AACTTGCTCAGGAGAACGCA |
| *Bai1 F* | TTGCTCCACTCCTGCTGTTAC |
| *Bai1 R* | GTAGCCGAAGAACTTTCCCTG |

Hprt, hypoxanthine guanine phosphoribosyl transferase; Col1a1, collagen type I alpha 1; Col1a2, collagen type I alpha 2; Col3a1, collagen, type III alpha 1; Tgfb1, transforming growth factor, beta 1; Acta2, actin alpha 2, smooth muscle, aorta; Spp1, secreted phosphoprotein 1; Timp1, TIMP metallopeptidase inhibitor 1; F4/80, adhesion G protein-coupled receptor E1; Tnfa, tumor necrosis factor-α ; Il1b, interleukin 1 beta; Mcp1, monocyte chemoattractant protein-1; Timd3, T-Cell Immunoglobulin And Mucin Domain-Containing Protein 3; Timd4, T-Cell Immunoglobulin And Mucin Domain-Containing Protein 4; MerTK, MER proto-oncogene tyrosine kinase; Axl, AXL receptor tyrosine kinase; Lrp1, LDL receptor related protein 1; Trem2, triggering receptor expressed on myeloid cells-2; Bai1, adhesion G protein-coupled receptor B1
